# Homozygous *TRPV4 mutation* causes congenital distal spinal muscular atrophy and arthrogryposis

**DOI:** 10.1101/402388

**Authors:** Jose Velilla, Michael Mario Marchetti, Agnes Toth-Petroczy, Claire Grosgogeat, Alexis H Bennett, Nikkola Carmichael, Elicia Estrella, Basil T. Darras, Brigham Genomic Medicine, Natasha Y Frank, Joel Krier, Rachelle Gaudet, Vandana A. Gupta

## Abstract

**Objective:** The objective of this study is to identify the genetic cause of disease in a congenital form of congenital spinal muscular atrophy and arthrogryposis (CSMAA).

**Methods:** A 2-year-old boy was diagnosed with arthrogryposis multiplex congenita, severe skeletal abnormalities, torticollis, vocal cord paralysis and diminished lower limb movement. Whole exome sequencing was performed on the proband and family members. *In silico* modeling of protein structure and heterologous protein expression and cytotoxicity assays were performed to validate pathogenicity of the identified variant.

**Results:** Whole exome sequencing revealed a homozygous mutation in the *TRPV4* gene (c.281C>T; p.S94L). The identification of a recessive mutation in *TRPV4* extends the spectrum of mutations in recessive forms of the *TRPV4*-associated disease. p.S94L and other previously identified TRPV4 variants in different protein domains were compared in structural modeling and functional studies. *In silico* structural modeling suggests that the p.S94L mutation is in the disordered N-terminal region proximal to important regulatory binding sites for phosphoinositides and for PACSIN3, which could lead to alterations in trafficking and/or channel sensitivity. Functional studies by western blot and immunohistochemical analysis show that p.S94L reduces TRPV4 protein stability because of increased cytotoxicity and therefore involves a gain-of-function mechanism.

**Conclusion:** This study identifies a novel homozygous mutation in *TRPV4* as a cause of the recessive form of congenital spinal muscular atrophy and arthrogryposis.

---

Hereditary neuropathies are a clinically and genetically heterogeneous group of diseases with an estimated prevalence of 1:2500. Clinical manifestations of hereditary neuropathies include slow progressive distal weakness and muscle wasting with or without sensory loss. Hereditary neuropathies are classified in to three broad categories on the basis of clinical phenotype. Charcot-Marie-Tooth (CMT) disease or hereditary motor and sensory neuropathy (HMSN) typically exhibits involvement of both motor and sensory systems, hereditary sensory and autonomic neuropathy (HSAN) involves sensory deficits and/or autonomic dysfunction, and distal hereditary motor neuropathy (dHMN) predominantly involves motor deficits. These groups can be further classified into many subtypes depending on electrophysiological criteria, pathological defects, mode of inheritance and molecular genetic defects. These diseases are genetically highly heterogeneous with mutations in at least 80 different genes associated with these subtypes^1, 2^. Despite this progress, 30-70% of people affected with neuropathies do not have a genetic diagnosis due to clinical and genetic heterogeneity. Some of disease symptoms can be controlled with generic drugs. However, the identification of the underlying genetic lesion is necessary for accurate disease prognosis, management and family planning, and may ultimately lead to personalized treatments.

Mutations in the *TRPV4* (transient receptor potential vanilloid 4) cation channel gene are a rare cause of dominant inherited axonal neuropathies and skeletal dysplasias ^3, 4^. *TRPV4* mutations underlie a wide spectrum of clinical presentation and are associated with dHMN, scapuloperoneal spinal muscular atrophy (SPSMA), congenital spinal muscular atrophy and arthrogryposis (CSMAA), autosomal dominant axonal CMT type 2C, and congenital distal spinal muscular atrophy (dSMA). *TRPV4*-related neuropathies are frequently associated with vocal cord paralysis and occasionally with sensorineural hearing loss. *TRPV4* mutations can also result in autosomal dominant skeletal dysplasias. Typically, different mutations in *TRPV4* gene are associated with either neuropathies or skeletal dysplasia; however, some patients exhibit both clinical phenotypes. So far, >20 different mutations in *TRPV4* have been identified in patients with neuropathies. However, functional validation of pathogenicity for many of these variants is still lacking, thus preventing a conclusive genetic diagnosis in these patients. To improve the genetic diagnosis of patients affected with neuropathies, we are using whole exome sequencing (WES) combined with functional validation studies to establish the pathogenicity of variants identified by WES. This work has identified a *TRPV4* homozygous mutation in a patient presented with distal hereditary motor neuropathy. Follow-up functional studies reveal distinct disease mechanisms for different TRPV4 variants and provide novel insights that may inform future therapeutic strategies for these patients.

## Materials and Methods

### Standard protocol approvals, registrations, and patient consents

The proband, both parents, and the unaffected sibling were enrolled and informed consent was obtained from participants in accord to an Institutional Review Board approved study at Boston Children’s Hospital and Brigham and Women’s Hospital.

### Whole-exome sequencing

DNA extraction from blood samples was performed by the Research Connection Biobank Core (Boston Children’s Hospital) using QIAmp DNA Mini kit (Qiagen). Whole exome sequencing was performed by the Yale Genome Center. DNA samples from the proband and parents were sent for whole exome sequencing. Samples were prepared as an Illumina sequencing library and enriched for exomic sequences using the Agilent V5 Sureselect kit. The captured libraries were sequenced using Illumina HiSeq 2000 Sequencers at Lab Corp. FASTQs generated from exome sequencing were filtered and aligned, and variants were filtered and annotated, as previously described^5^. In short, first we apply agnostic filtering, and we use pedigree-based inheritance mode filtering for *de novo* and recessive variants, as well as filter for rare variants only, based on large population databases (gnomAD, gnomad.broadinstitute.org). In the second step, we apply several knowledge-based filters on the genes (e.g. known functional and disease associations, expression level data, model organism data) and on the variants (e.g. evolutionary conservation, structural constraints etc.). Finally, the variants were prioritized during a crowdsourcing case conference of interdisciplinary audience.^5^To validate the *TRPV4* variant, PCR products for the proband, unaffected sibling and parents were analyzed by standard Sanger sequencing (Dana-Farber/Harvard Cancer Center DNA Sequencing Facility).

### *In silico* modeling of TRPV4 mutations

TRPV4 mutations were mapped onto the cryoEM structure of *Xenopus tropicalis* TRPV4 (PDB ID: 6bjj)^6^ after aligning the XtTRPV4 and human TRPV4 sequences using Clustal Omega. Figure 2 was generated using PyMOL (Schrodinger).

**Figure 2.**
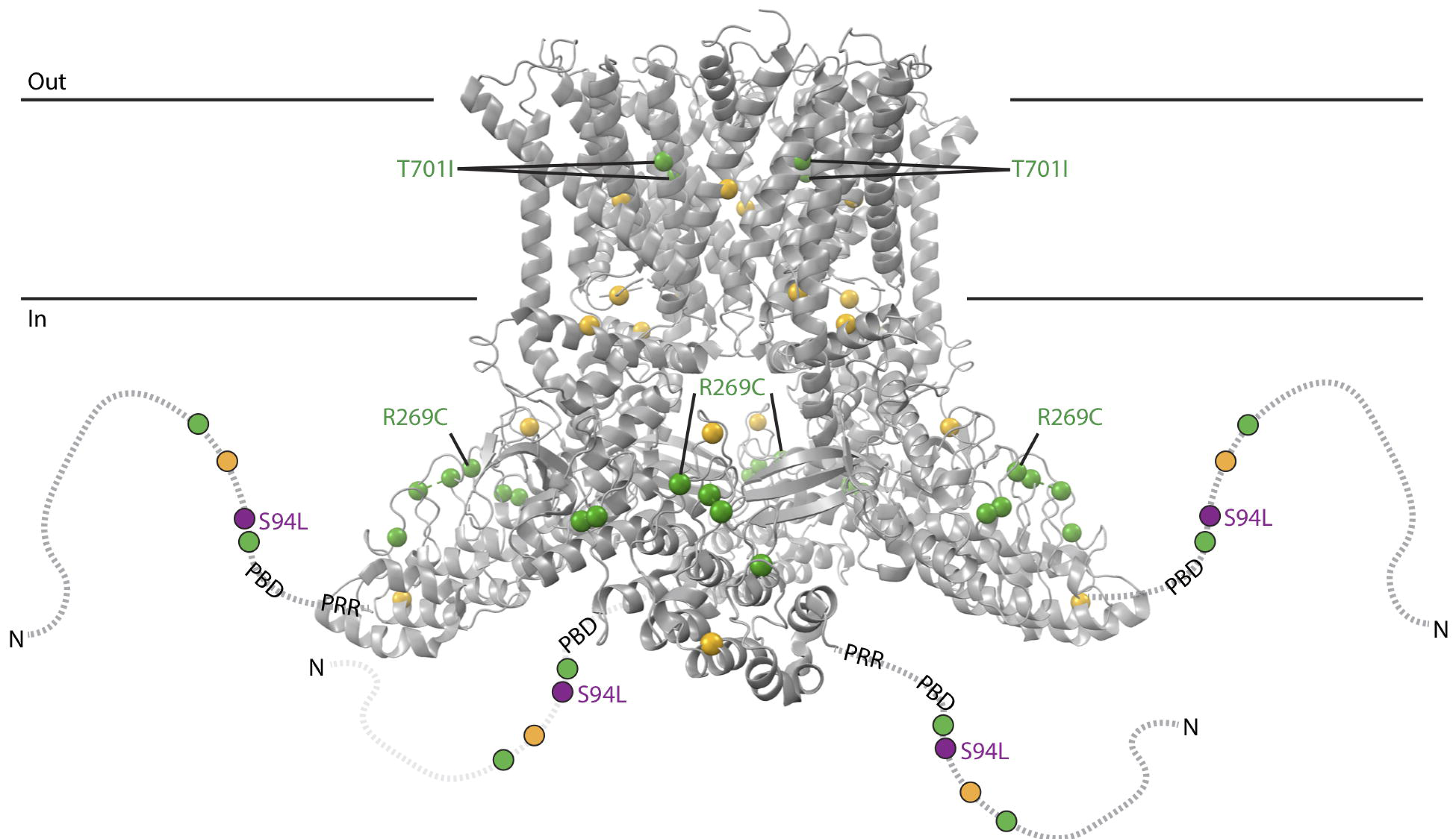
Structural model based on the cryoEM structure of *Xenopus tropicalis* TRPV4 (PDB ID: 6bjj). The structure, corresponding to residues 148-788 (human numbering) does not include disordered N- and C-terminal regions. The N-terminal region is schematized as a dotted line for each subunit, with the phosphoinositide-binding domain (PBD; residues 121-125 in human) and proline-rich region (PRR, residues 135-144) indicated. Residue positions for neuropathy-causing mutations (D62N, P97R, R186Q, R232C/S, R237G/L, R269C/H, R315W, R316C/H, T701I) and disease mutations with mixed phenotypes (G78W, A217S, E278K, S542Y, V620Y, T740I) are green and yellow, respectively. Position of p.S94L is indicated in purple.

### Cloning of mutant constructs

To generate *TRPV4* mutant plasmids, p.S94L, p.R315W and & p.T701I variants were incorporated into *TRPV4* cDNA in plasmid pcDNA3.1-TRPV4-FLAG using Q5 site-directed mutagenesis kit (New England Biolabs). The mutagenesis primer sequences were: S94L forward: 5’-TAT GAG TCC TTG GTG GTG CCT-3’, S94L reverse: 5’-TAG GGT GGA CTC CAG CAG-3’, R315W forward: 5’-GGC GGA CAT GTG GCG CCA GGA-3’, R315W reverse: 5’-TTC TTG TGG GGG TTC TCC GTC AGGT-3’, T701I forward: 5’-CTG CTG GTG ATC TAC ATC ATC-3’, T701I reverse: 5’-GAT GAT GAA GAC CAC GGG-3’. The full coding sequences were confirmed using Sanger sequencing.

### Cytotoxicity Assay

Human HEK293T cells were cultured in DMEM supplemented with 10% fetal bovine serum. Cells were transfected with the respective TRPV4 or empty vector using Lipofectamine 3000 (Thermo Fisher Scientific) and cultured in the presence or absence of HC-067047 (5 μM). Cell death analysis was performed 24 hours post-transfection using the Cytotoxicity Detection Kit (Roche Diagnostics, Indianapolis, IN) according to the manufacturer’s instructions. Cytotoxicity was calculated by subtracting the absorbance of the background control (media from empty vector condition) from the absorbance of the experimental samples (supernatant from cells transfected with WT or mutant plasmid).

### Western Blot

HEK293 cells were transfected with the respective vector using polyethylenimine (PEI to DNA ratio of 3:1). Cells were incubated for 12 hours at 37°C, then shifted to 33°C, and harvested 48 hours post-transfection. Western blots were performed according to standard protocols, using 1:2000 M2 anti-FLAG (Sigma, F-3165) to detect TRPV4, 1:10000 anti-GAPDH (Abcam, 8245), and 1:1000 alkaline phosphatase conjugated anti-mouse secondary antibody (Sigma, A3562). Membranes were developed using the 1-Step NBT/BCIP reagent (ThermoFisher Scientific, 34042). Densitometric analysis was performed with imageJ (https://imagej.nih.gov/ij/).

### Immunofluorescence Analysis

For immunofluorescence, HEK293T cells were plated in 8-chambered slides and transfections were performed with lipofectamine 3000 (Thermo Fisher Scientific). Twenty-four hours after transfection, cells were fixed in 4% paraformaldehyde and immunofluorescence was performed as described previously^16^. M2 anti-FLAG antibody (Sigma, 1:250) and sodium potassium ATPase antibody (Abcam 76020,1:100). Nuclear staining was done using DAPI (Biolegend, 422801).

## Results

### Clinical Presentation

The male proband was born to unrelated healthy parents of Puerto Rican descent after 37 weeks of gestation (Fig. 1A). He has a healthy sister. The proband was delivered by Caesarean section due to breech position and arthrogryposis multiplex congenita on ultra-sound. At birth, the proband exhibited a right clubfoot, a left congenital vertical talus, and bilateral flexion deformities of the knees. X-rays showed dislocated hips with wide proximal femurs. Ultrasound revealed that patient exhibited dysplastic acetabuli on both sides. An ultrasound of the spine was done which showed that spinal cord was at the L3 level. Bilateral flexible laryngoscopy was performed through left naris that showed nonobstructive adenoid hypertrophy, normal tongue base, structurally normal larynx with no evidence of laryngomalacia. Vocal cords were clearly visible and appeared to be immobile bilaterally in the paramedian position. The proband also exhibited inspiratory stridor. Examination of right and left tympanic membranes showed an evidence of middle ear effusions bilaterally and normal external ear and external auditory canals. Visual reinforced audiometry showed a moderate conductive hearing loss for at least one ear at 500 and 2000 Hz. Tympanograms were flat bilaterally consistent with clinical findings.

**Figure 1.**
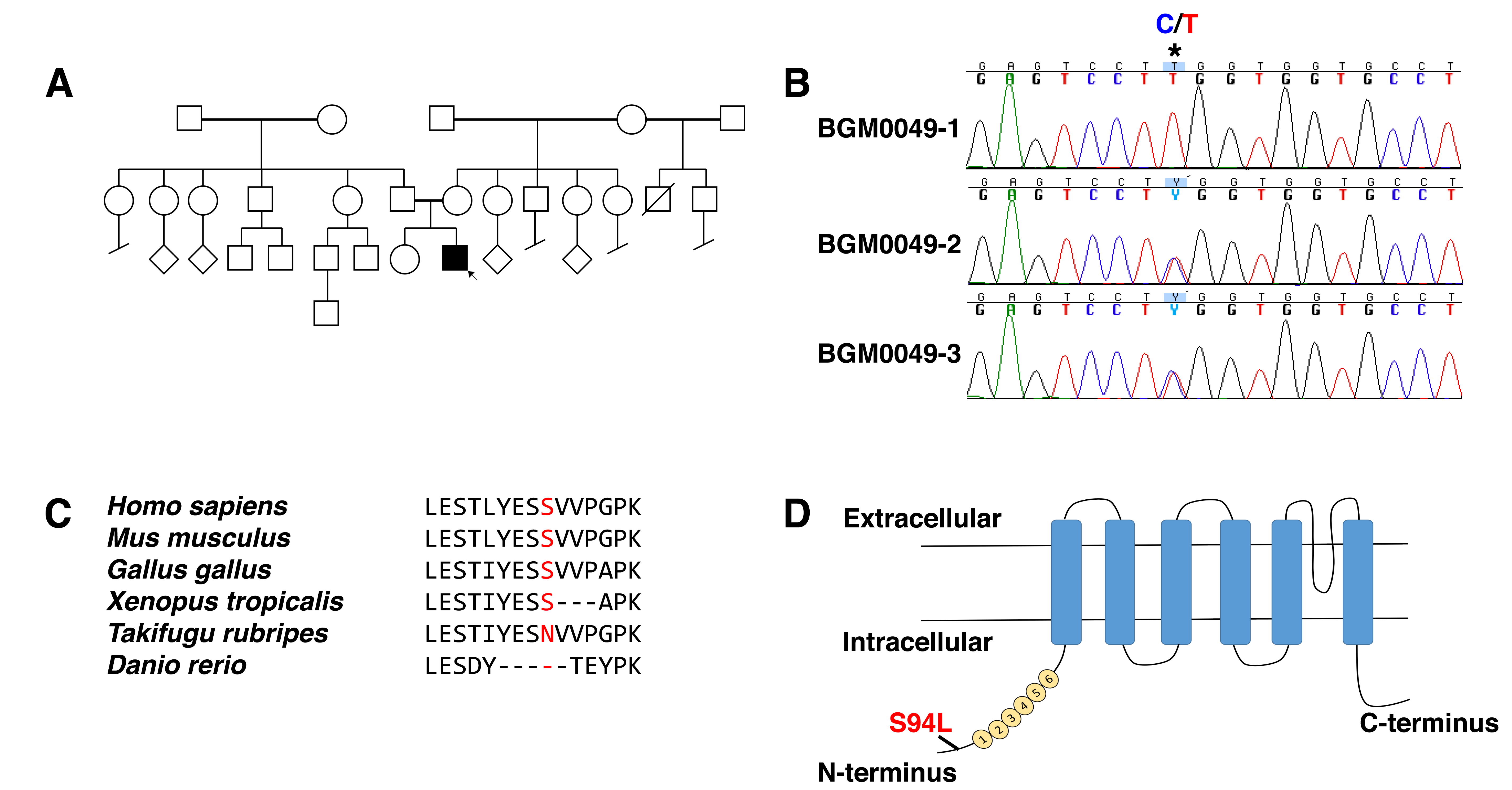
(A) Pedigree of family of affected patient; proband is indicated by the arrow (B) Sanger sequencing pherograms show homozygosity for c.281C>T in *TRPV4* in the proband and heterozygosity in both parents. (C) Protein sequence alignment of TRPV4 orthologs showing conservation in the region including p.S94L in higher vertebrates. (D) Schematic diagram of the TRPV4 protein demonstrating localization of p.S94L on the N-terminal intercellular region. Numbers 1-6 correspond to the six ankyrin repeats.

At age 2 years, the proband could sit independently and walk on his knees. However, he was unable to stand or walk independently. Except for lower limb movement, the proband has normal speech, fine motor and gross motor skills in his upper limbs, normal hearing and visual development and exhibit a static condition.

Electromyography (EMG) examination revealed normal right rural sensory response and normal left ulnar-DV sensory response. Normal left tibial-AH and median-APB motor response for the age was also noted. Concentric needle examination of selected muscles showed fibrillation potentials in the right vastus laterals muscle with late and fast firing motor unit potentials in left deltoid, right tibialis anterior, and right iliopsoas muscles. The electrophysiological findings are suggestive of generalized motor axonopathy with coexisting denervation and reinnervation changes.

### Whole Exome Sequencing (WES)

Whole exome sequencing was performed on the proband and parents. The presence of a single affected male individual in this family is consistent with autosomal recessive inheritance, X-linked inheritance or with a dominant *de novo* mutation. Therefore, pedigree and population based filtering was performed to identify potentially damaging homozygous, X-linked, compound heterozygous and *de novo* variants. We only considered rare coding and splice variants that have <0.1-1 % allele in general healthy population in the gnomAD database (gnomad.broadinstitute.org) depending on dominant or recessive inheritance mode, that is consistent with the prevalence of the disease. In total, 15 potential genes carried a variant (Table e-1), that were further prioritized using our knowledge-based and crowdsourcing pipeline recently reviewed in Haghighi et al ^5^. The top candidate was a homozygous missense variant in *TRPV4* c.281C>T; (p.S94L). Sanger sequencing confirmed the homozygosity of this variant in proband (BGM0049-1) and heterozygous inheritance from both parents (BGM0049-2, -3) (Fig. 1B). The unaffected sister was found to be homozygous for the normal variant (not shown). The other 14 variants/genes could be ruled out by having low population constraints, known unrelated disease associations, unrelated protein functions and lack of expression in relevant tissues (Table e-1), corroborating the potential causal effect of the *TRPV4* variant. This missense variant is highly conserved in most of the vertebrate species and is localized to N-terminus of the TRPV4 protein (Fig. 1C-D).

### Structural Model

A recent cryoelectron microscopy (cryoEM) structure of *Xenopus tropicalis* TRPV4 was used to model the position of p.S94L and other previously described neuropathy-causing TRPV4 mutations (Fig. 2)^6^. Consistent with previous proteolysis protection^7^ and nuclear magnetic resonance studies^8^ the N-terminal region – up to residue 147 (human numbering), thus including p.S94 – was disordered and not modeled in the cryoEM structure. However, p.S94L is located N-terminal to two important regulatory regions: a phosphoinositide-binding domain (PBD; residues 121-125 in humans) that interacts with PIP_2_^7^ and a proline-rich region (PRR; residues 135-144 in humans) that interacts with the SH3 domain of PACSIN3^9^. PACSIN3 enhances TRPV4 trafficking to the cellular membrane. PIP_2_ binding enhances TRPV4 responses to several stimuli whereas PACSIN3 decreases responses to these same stimuli via an antagonistic interaction between two key regulators, PIP_2_ and PACSIN3, that is likely mediated through PACSIN3-mediated structural rearrangements of the PRR^7, 10^. Therefore, our structural modeling suggests that the p.S94L mutation could potentially affect the regulation and/or sensitivity of TRPV4.

### Functional Modeling of TRPV4 Variants

Functional studies have shown a gain-of-function mechanism for previously known dominant mutations in *TRPV4*^11^. To validate the pathogenicity of the p.S94L variant, functional studies were performed by analyzing the stability and localization of the mutant protein. To compare the magnitude of the functional deficits, known neuropathy-causing p.R269C and p.R315W TRPV4 mutants were analyzed in parallel. In addition, a mutation p.T701I has been reported as a cause of neuropathy. p.T701I is localized to the transmembrane domain S6 helix (Fig. 2), near the vanilloid-binding site in the homologous ion channel TRPV1.^12^ Currently, there are no functional studies establishing the pathogenicity of this variant in human diseases or TRPV4 protein functions. Therefore, we investigated the effect of p.S94L, p.R269C, p.R314W and p.T701I on the stability and subcellular localization of TRPV4. To investigate the effect of different variants on protein stability, TRPV4-WT and mutant constructs were transiently expressed in HEK293T cells and expression was analyzed by western blot (Fig. 3). Protein analysis by western blot showed reduced level of p.S94L protein as well as reduced levels of p.R269C and p.R315W. In contrast, p.T701I exhibited levels of protein expression similar to those of wild-type TRPV4 (Fig. 3A-B). As previous studies have reported cytotoxic effects of mutant TRPV4 proteins, cytotoxicity analysis was performed in HEK293T cells transfected with control and mutant plasmids. Quantification showed significantly higher levels of cell death in the p.S94L transfected cells in comparison to wild-type TRPV4 transfected cells. Similarly, p.R269C and p.R315W also resulted in the increased cytotoxicity. Interestingly, no changes in cytotoxicity were observed in p.T701I transfected cells in comparison to wild-type TRPV transfected cells. HEK293T cells transfected with control and mutant plasmids were also cultured in the presence of a TRPV4 channel antagonist (HC-067047) that reversed the cytotoxicity effects observed in p. S94L, p.R269C and p.315W transfected cells (Fig. 3C). Presence of HC-067047 also resulted in an increased expression of p.S94L, p.R269C and p.R315W, however, no expression changes were detected for p.701I protein (Fig. 3B) To examine the sub-cellular localization of p.S94L and other mutants, immunofluorescence analyses were performed (Fig. 4). Wild-type TRPV4 primarily localized to the plasma membrane as has been reported for endogenous TRPV4 protein. p.S94L and previously reported p.R269C and p.R315W exhibited localization to the plasma membrane. Interestingly, the p.T701I mutant protein showed a perinuclear localization pattern, and in contrast to wild-type protein, no protein was detected on the plasma membrane (Fig. 4).

**Figure 3:**
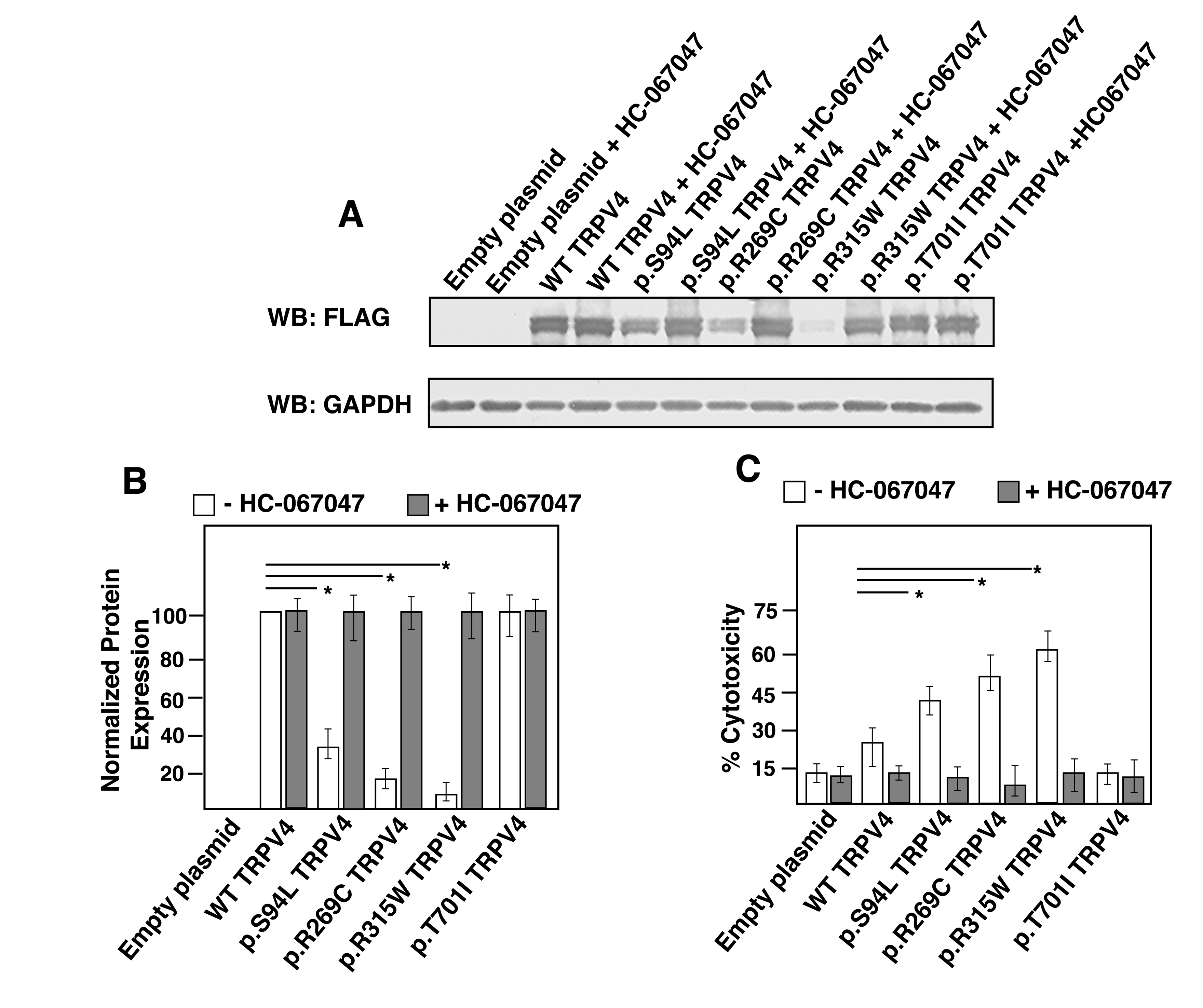
(A-B) Western blot from HEK293T cells transiently expressing TRPV4-FLAG, p.S94L-FLAG, p.R269C-FLAG, p.R315W-FLAG, p.T701I -FLAG or empty vector collected 48 hours after transfection and incubation in the absence or presence of HC-067047 (5 μM). Cell lysates were probed with antibodies against FLAG tag to detect transfected wild-type and mutant TRPV4 proteins, and GAPDH as the loading control. (C) Cytotoxicity analysis in cells transfected with control or mutant *TRPV4* plasmids. Differences in the protein levels or cytotoxicity were considered significant with * *p*<0.05.

**Figure 4:**
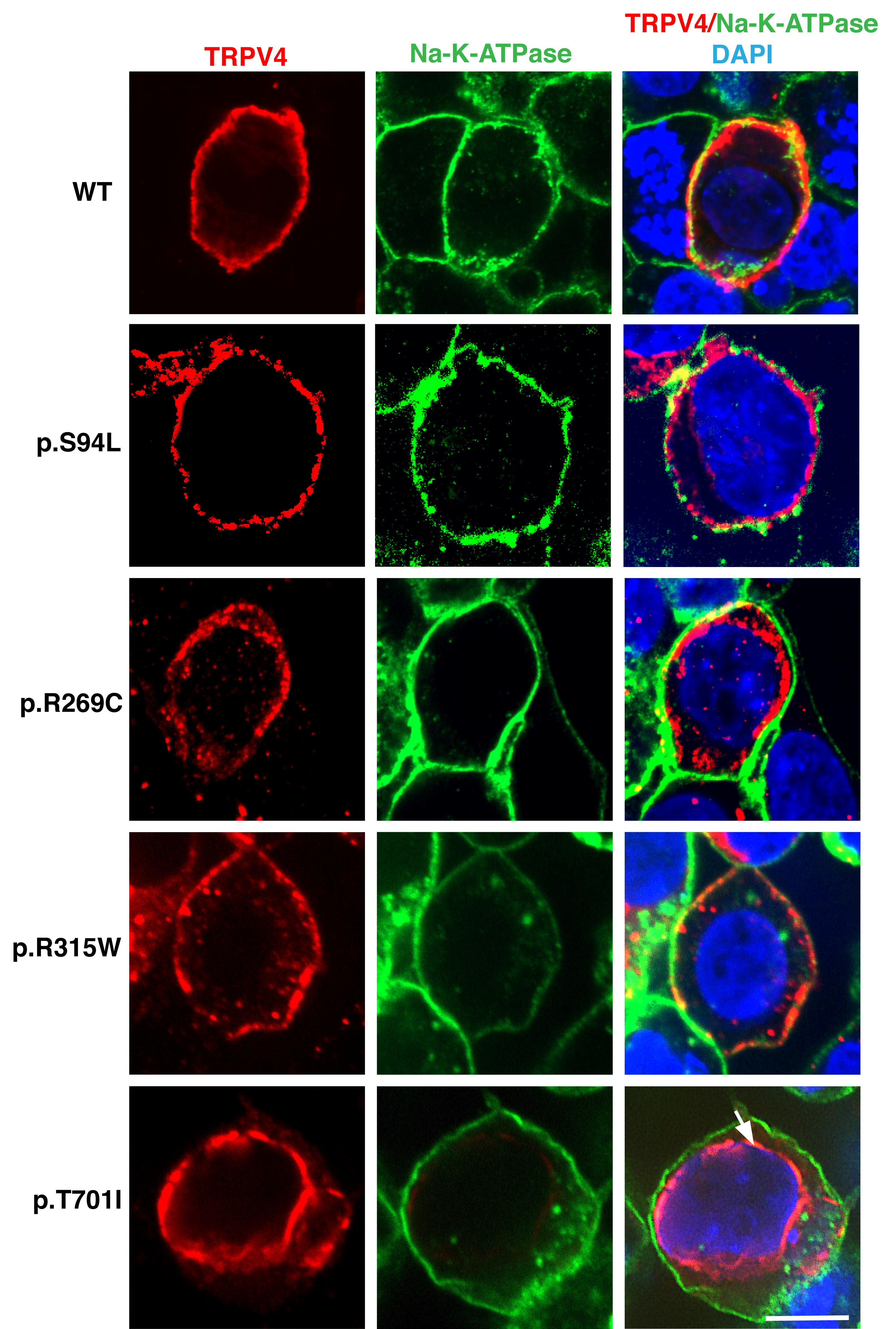
Representative images of HEK293T cells transiently expressing TRPV4-Flag, p.S94L-Flag, p.R269C-Flag, p.R315W-Flag, p.T701I proteins. Immunofluorescence was performed with anti-Flag tag antibody to detect TRPV4-Flag proteins and Na-K-ATPase to label the plasma membrane. Nuclei were stained with DAPI. Scale bar = 20μM.

## Discussion

The distal hereditary motor neuropathies (dHMN) are a genetically heterogeneous group of diseases characterized by distal lower-motor-neuron weakness. This is in contrast to CMT disease and the hereditary sensory neuropathies where sensory involvement forms a major component of the disease. Many forms of dHMN can exhibit a minor sensory component and, there is an overlap between axonal forms of CMT (CMT2) and dHMN. Autosomal dominant mutations in *TRPV4* are associated with diverse clinical presentations including dHMN, SPSMA, CSMAA, and autosomal dominant axonal CMT type 2C (CMT2C) disease. *TRPV4* mutations also result in several forms of autosomal dominant skeletal dysplasias. The proband exhibited arthrogryopsis associated with distal muscle weakness with normal sensory response and therefore, clinically diagnosed as congenital spinal muscular atrophy and arthrogryposis (CSMAA), a subtype of dHMN. Several functional studies have demonstrated a gain-of-function mechanism for mutant TRPV4-mediated neuropathies^11^. Mutant TRPV4 proteins frequently exhibit normal localization; however, they demonstrate increased calcium channel activity at both basal and activated levels. This results in increased intracellular calcium concentration leading to cytotoxicity.

Following a combination of whole exome sequencing and functional analysis, we identified a homozygous missense variant, p.S94L, in TRPV4 as a cause of a severe form of distal hereditary motor neuropathy. This serine residue is highly conserved in mammals and only 6 heterozygous and no homozygous individuals are reported in a control population of 246210 people (Genome Aggregation Database, gnomAD), implicating this as a crucial amino acid. Here, we demonstrate that the TRPV4 p.S94L variant resulted in increased cytotoxicity, as previously reported for other neuropathy-causing pathogenic TRPV4 variants. A limitation of this work is that these studies were performed in HEK293T cells rather than neuronal cells. However, HEK293T cells have commonly been used to determine aberrant channel function for dominant *TRPV4* mutations, allowing comparisons of our results with previous work^3^. Furthermore, p.S94L mutation resulted in reduced amounts of the mutant protein that were rescued by the TRPV4 channel antagonist as previously observed in neuronal cells^11^. Previous work has also identified reduced stability of many dominant variants in the intracellular ankyrin-repeat domain of TRPV4^13^. A recent study also identified biallelic heterozygous mutations located in the C-terminus of the TRPV4 channel in two siblings with neuropathy associated with severe intellectual disability^14^. These mutant proteins demonstrated normal membrane localization; however, they showed reduced channel function. The severe phenotype in the proband is consistent with the observed cytotoxicity of the p.S94L variant in the HEK293T cells.

The p.S94L mutation is located in the N-terminal intracellular region upstream of the ankyrin repeat domain. Previous studies have shown that dominant mutations in this region result in neuropathy in affected individuals^15^. In comparison to previously known TRPV4 variants such as p.R269C and p.R315W, higher levels of p.S94L TRPV4 protein were detected in transfected HEK293T cells (Fig. 3 A-B). Similarly, we observed a lower level of cytotoxicity in p.S94L transfected cells in comparison with p.R269C and p.R315W transfected cells (Fig. 3C). Furthermore, the facts that the proband’s parents who are heterozygous for the *TRPV4* p.S94L variant did not exhibit any phenotype, and that six heterozygous individuals are reported in the gnomAD database, suggest reduced penetrance of this variant in the heterozygous state. Nonetheless, many heterozygous*TRPV4* variants exhibit reduced penetrance in adult carriers and for this reason, the parents of the proband were counseled to routine follow up in neurology clinic. Interestingly, most human disease-causing variants studied in this work exhibited a reduced protein stability, whereas the p.T701I variant altered the cellular localization of TRPV4 protein, from plasma membrane to the perinuclear area. This mis-localization may result in altered calcium channel function in affected patients. Together, these results demonstrate that different mutations in *TRPV4* affect the channel function by different mechanisms. Understanding the functional alterations in mutant TRPV4 channels is essential not only for determining the pathogenicity of novel variants, but also for the design of specific therapies as both agonists and antagonists of TRPV4 channels are available.

## Supporting information

## Data Availability

The research material supporting this publication can be accessed by contacting the corresponding author at.

## Acknowledgement

This work was supported by K01 AR062601 (VAG) and the Eleanor, Miles Shore Fellowship for Scholars in Medicine and Brigham and Women’s Hospital Career Development Award (VAG), a Brigham Biomedical Research Institute Director’s Transformative Award (Brigham Genomic Medicine), and the American Heart Association 16GRNT27250119 (RG).

## Appendix I

| Name | Location | Role | Contribution |
| --- | --- | --- | --- |
| Jose Velilla. BS | Harvard University | Author | Major role in the acquisition of data |
| Michael M Marchetti, BS | Brigham and Women's Hospital | Author | Major role in the acquisition of data |
| Agnes Toth-Petroczy, PhD | Brigham and Women's Hospital | Author | Interpreted the data; revised the manuscript for intellectual content |
| Claire Grogogeat | Brigham and Women's Hospital | Author | Major role in the acquisition of data |
| Alexis H. Bennett, MS | Brigham and Women's Hospital | Author | Major role in the acquisition of data |
| Nikkola Carmichael, MS | Brigham and Women's Hospital | Author | Interpreted the data; revised the manuscript for intellectual content |
| Elicia Estrella, MS | Boston Children's Hospital | Author | Design and conceptualized study; analyzed the data; drafted the manuscript for intellectual content |
| Basil T. Darras, MD | Boston Children's Hospital | Author | Design and conceptualized study; analyzed the data; drafted the manuscript for intellectual content |
| Natasha Y. Frank, MD | Brigham and Women's Hospital | Author | Interpreted the data; revised the manuscript for intellectual content |
| Joel Krier, MD | Brigham and Women's Hospital | Author | Interpreted the data; revised the manuscript for intellectual content |
| Rachelle Gaudet, PhD | Harvard University | Author | Design and conceptualized study; Interpreted the data; revised the manuscript for intellectual content |
| Vandana A. Gupta, PhD | Brigham and Women's Hospital | Author | Design and conceptualized study; analyzed the data; drafted the manuscript for intellectual content |

## Figure Short Titles

Figure 1. Identification of TRPV4 homozygous mutation

Figure 2. Structure modeling of TRPV4 mutations

Figure 3. Impact of TRPV4 mutations on protein stability and cytotoxicity

Figure 4. Subcellular localization of mutant TRPV4 proteins

---

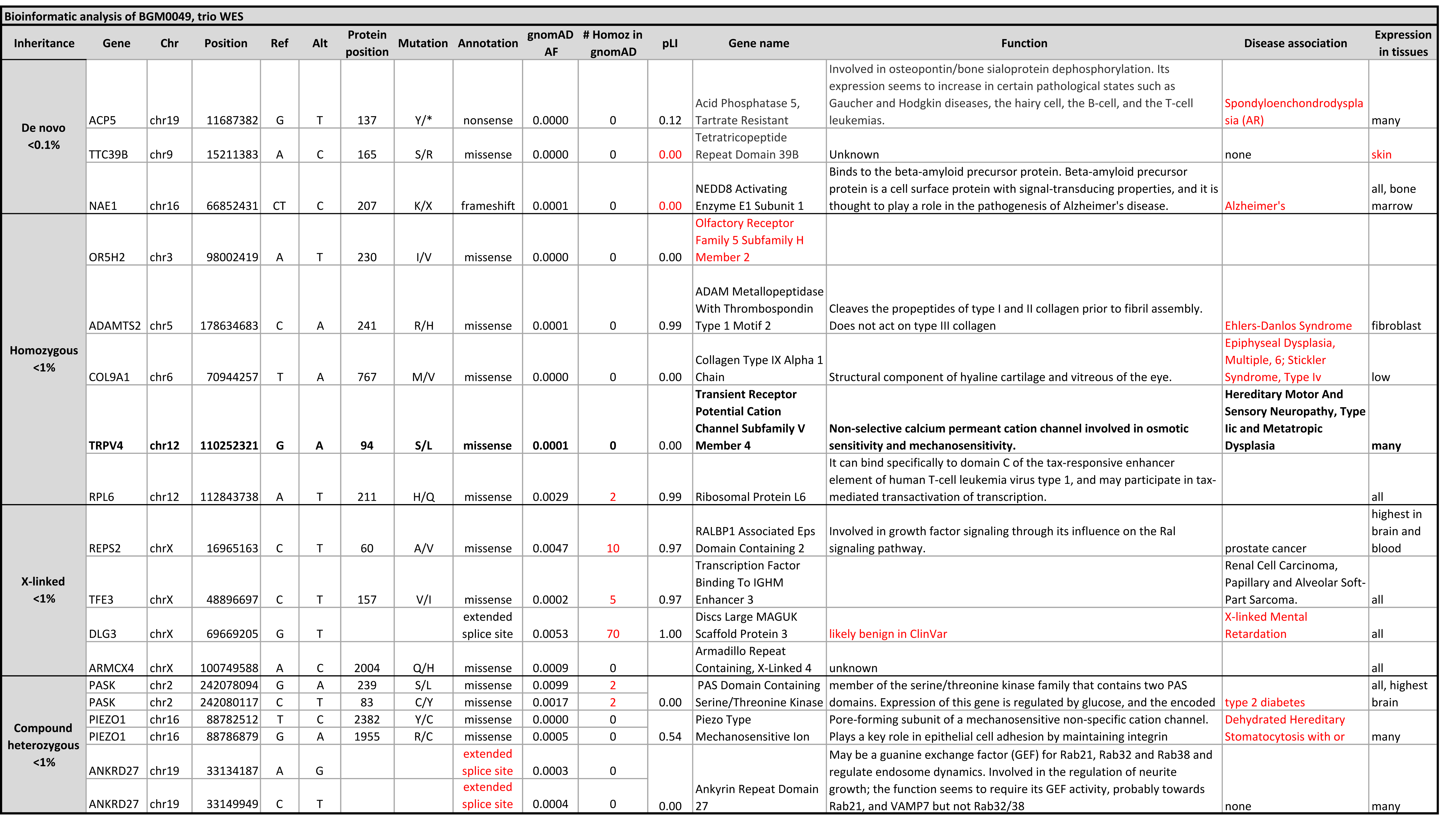

## Glossary

dHMN: distal hereditary motor neuropathy
HMSN: hereditary motor and sensory neuropathy
HSAN: hereditary sensory and autonomic neuropathy
SPSMA: scapuloperoneal spinal muscular atrophy
CSMAA: congenital spinal muscular atrophy and arthrogryposis
CMT: Charcot-Marie-Tooth diseases
PBD: phosphoinositide-binding domain
PRR: proline-rich region
PIP_2_: phosphatidylinositol 4,5-biphosphate
FDAB: Familial digital arthropathy-brachydactyly

## Notes

#### Summary of Updates

Figures 3 and 4 are revised

